# Rap1 Activation Protects Against Fatty Liver and Non-Alcoholic Steatohepatitis Development

**DOI:** 10.1101/2023.10.24.563728

**Authors:** Heena Agarwal, Yating Wang, Lale Ozcan

**Author notes:** **Corresponding author:** Lale Ozcan MD, 630 West 168^th^ Street, Black Building: 901D, New York, NY 10032, USA.

## Abstract

We previously demonstrated that hepatic activation of a small G protein of the Ras family, Rap1a, is suppressed in obesity, which results in increased hepatic glucose production and glucose intolerance in obese mice. Here, we show that Rap1a inhibition in obese mice liver also results in fatty liver formation, which is characteristic of the diabetic liver. Specifically, we report that Rap1a activity is decreased in the livers of patients with non-alcoholic steatohepatitis (NASH) and mouse models of non-alcoholic fatty liver disease (NAFLD) and NASH. Restoring hepatic Rap1a activity by overexpressing a constitutively active mutant form of Rap1a lowered the mature, processed form of lipogenic transcription factor, Srebp1, without an effect on the unprocessed Srebp1 and suppressed hepatic TG accumulation, whereas liver Rap1a deficiency increased Srebp1 processing and exacerbated steatosis. Mechanistically, we show that mTORC1, which promotes Srebp1 cleavage, is hyperactivated upon Rap1a deficiency despite disturbed insulin signaling. In proof-of-principle studies, we found that treatment of obese mice with a small molecule activator of Rap1a (8-pCPT) or inhibiting Rap1a’s endogenous inhibitor, Rap1Gap, recapitulated our hepatic gain-of-function model and resulted in improved hepatic steatosis and lowered lipogenic genes. Thus, hepatic Rap1a serves as a signaling molecule that suppresses both hepatic gluconeogenesis and steatosis, and inhibition of its activity in the liver contributes to the pathogenesis of glucose intolerance and NAFLD/NASH development.

## Introduction

Type 2 diabetes (T2D) and its complications constitute a significant public health problem worldwide and are an important cause of morbidity and mortality.^1^ While current treatment options for T2D have beneficial impact on the disease and some of its complications, there is a great need for novel therapeutic approaches to improve long-term glycemic control and T2D-related complications. Hepatic glucose production (HGP) is crucial for systemic glucose homeostasis and an increase in HGP is a key feature of T2D.^2,3^ Several factors, including the availability of substrates and an imbalance between glucagon and insulin action contribute to this process.^4^ On the other hand, insulin induces the expression of hepatic sterol regulatory element-binding transcription factor (Srebp1), which upregulates the genes involved in de novo lipogenesis (DNL).^5^ Insulin also suppresses adipose tissue lipolysis and lowers the amount of fatty acyl-CoA available for esterification into triglycerides (TG) in the liver. In insulin-resistant individuals, insulin fails to inhibit HGP owing to defects in insulin receptor signaling; however, its ability to stimulate hepatic lipid synthesis and storage is unimpaired.^6,7^ This selective insulin resistance results in the characteristic hyperglycemia and non-alcoholic fatty liver disease (NAFLD) of T2D.^8^ In this context, identification of molecular targets that uncouple insulin’s effects on hepatic lipogenesis from lowering HGP has a tremendous potential to lead to novel and more specific anti-diabetic drugs.

20% of people with NAFLD progress to a more severe form of liver disease referred to as non-alcoholic steatohepatitis (NASH).^9^ NASH is characterized by steatosis, hepatocyte injury, inflammation, and most importantly, fibrosis, which is the main determinant of cirrhosis and hepatocellular carcinoma.^10,11^ Enhanced lipid flux and DNL mediated increases in lipotoxic stress results in hepatocyte injury, which in turn initiates an immune and fibrogenic response driven primarily by Kupffer and hepatic stellate cells, respectively.^11^ There are currently no FDA approved therapies for NASH. Therefore, investigation of molecular mechanisms by which steatosis progresses to NASH-induced fibrosis could result in new therapeutic targets.

We previously reported that Rap1a activity is lowered in obese mice liver, which results in increased HGP and hyperglycemia via suppressing the insulin mediated Akt activation.^12^ We also uncovered that the widely used cholesterol lowering drugs, statins, inhibit Rap1a prenylation and activity in isolated hepatocytes and human liver, which contributes to statins’ hyperglycemic effects.^12^ Here we show that Rap1a inhibition also results in increased lipogenesis and hepatic TG accumulation, whereas Rap1a activation has the opposite effects. Mechanistically, we demonstrate that Rap1a contributes to lipogenesis homeostasis through mTORC1 mediated Srebp1 regulation. Further, we report that Rap1a activation is suppressed in a mouse model of NASH and in the livers of patients with NASH and restoring its activity lowers plasma ALT and liver fibrogenic gene expression. Thus, our results identify hepatic Rap1a as a mechanistic node in obesity-related hyperglycemia and NAFLD/NASH pathogenesis and a bona fide therapeutic target.

## Results

### Restoring hepatic Rap1a activity lowers steatosis and hepatic lipogenic gene expression whereas Rap1a inhibition increases them in obese mice

Rap1a activity is suppressed in mouse models of obesity with fatty liver formation and this in turn contributes to increased gluconeogenesis and HGP.^12^ As the rates of gluconeogenesis and the extent of liver fat in NAFLD patients are closely related, we became interested in the role of Rap1a in liver TG accumulation. When Rap1a activity was restored via treatment of *db/db* mice with adeno-associated virus 8 (AAV8) that expresses CA-Rap1a driven by the hepatocyte-specific thyroxine binding globulin (TBG) promoter (AAV8-TBG-CA-Rap1a), we observed that CA-Rap1a expressing *db/db* mice had fewer lipid droplets (**Figure 1A-1B**) and decreased TG content (**Figure 1C**) without changes in body weight, food intake or plasma insulin levels (data not shown). To determine the mechanism of protection from hepatic TG accumulation in the TBG-CA-Rap1a-treated *db/db* mice, we examined lipid mobilization and metabolism pathways.^13^ We found that serum levels of TGs and hepatic TG output as assessed with Poloxamer-407 injection were not increased, in fact they were decreased, by TBG-CA-Rap1a treatment (not shown). Consistently, mRNA levels of proteins involved in TG mobilization, such as *Mttp, Dgat1*, and *Dgat2*, were not significantly different between TBG-CA-Rap1a treated vs control *db/db* mice (not shown). Furthermore, hepatic expression of fatty acid oxidation genes, including *Ppara, Acox* and *Cpt1a* were not increased in TBG-CA-Rap1a expressing mice (not shown). Collectively, these data suggest that neither an increase in hepatic lipid output nor stimulation of lipid oxidation are responsible for the protection from steatosis exhibited by hepatic Rap1a activation. We next turned our attention to lipogenesis and found decreased expression of sterol regulatory element–binding protein 1 (*Srebf1*) and lipogenic genes such as fatty acid synthase (*Fas*) and stearoyl-CoA desaturase (*Scd1*) in TBG-CA-Rap1a treated mice compared with controls (**Figure 1D**). Other regulators of lipogenic genes, including *Ppparg* signaling,^14^ were unaffected in TBG-CA-Rap1a vs control *db/db* mice (not shown). Consistently, TBG-CA-Rap1a treated mice had lower plasma ALT levels, suggesting less hepatocellular injury (**Figure 1E**). Conversely, we found that when Rap1a was deleted specifically in hepatocytes in diet-induced obese (DIO) *Rap1a*^fl/fl^ mice via injecting the AAV-TBG-Cre (L-Rap1a KO), the mice had increased hepatic TG and elevated plasma ALT levels as compared to AAV-TBG-Gfp injected control *Rap1a*^fl/fl^ mice (**Figure 1F-1G**). Thus, hepatic Rap1a activity restoration in obese mice lowers liver TG accumulation and lipogenic gene expression without a change in hepatic lipid output or lipid oxidation pathways and Rap1a deficiency has the opposite results.

**Figure 1.**
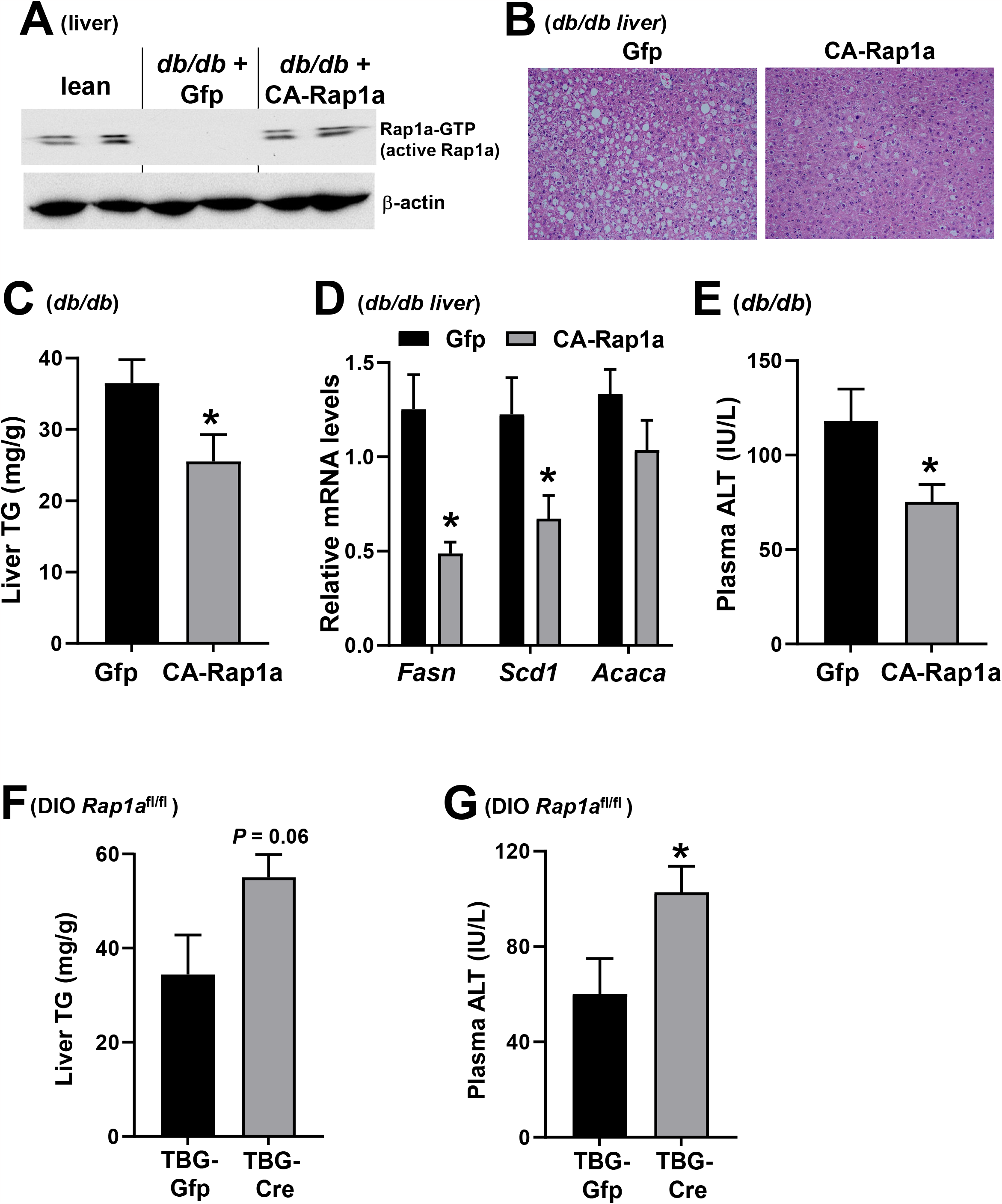
Restoring Hepatic Rap1a activity lowers steatosis and hepatic lipogenic gene expression whereas Rap1a inhibition increases them in obese mice. (**A**) Hepatic GTP-Rap1a (active Rap1a) levels in lean, TBG-Gfp treated *db/db* (control) and TBG-CA-Rap1a treated *db/db* mice. (**B**) Liver sections from TBG-Gfp and TBG-CA-Rap1a treated *db/db* mice were stained with H&E, and hepatic triglyceride (TG) content (**C**), liver lipogenic gene expression (**D**) and plasma ALT levels were analyzed (**E**) (n= 4-5; *p < 0.05). (**F-G**) Liver triglyceride (TG) content (F) and plasma ALT levels (G) were assayed in *Rap1a*^fl/fl^ DIO mice treated with AAV8-TBG-Cre or AAV8-TBG-Gfp (control) (n= 4-5 mice/ group, *p < 0.05).

### Rap1a activation via 8-pCPT or sh-Rap1Gap treatments lowers steatosis in DIO mice

We next asked whether activating Rap1a pharmacologically could have metabolic benefits and protect mice against hepatosteatosis. We used the specific Epac activator, 8-pCPT-2’-O-Me-cAMP (8-pCPT), which is a cAMP analog that confers specific Epac binding and robustly activates Rap1a.^15^ We and others have shown that Rap1 activation using 8-pCPT protects against ischemia-reperfusion injury-induced renal failure and asthmatic airway inflammation and lowers the proatherogenic PCSK9 levels.^16-18^ We observed that when DIO mice were treated with 8-pCPT for 2 weeks by daily injecting it at a dose of 1.5 mg/kg body wt, the mice had lower hepatic TG accumulation and decreased plasma ALT levels (**Figure 2A-2B**). Similar to Rap1a restoration, 8-pCPT treatment lowered hepatic expression of lipogenic genes (**Figure 2C**) in obese mice without a change in lipid oxidation (not shown).

**Figure 2.**
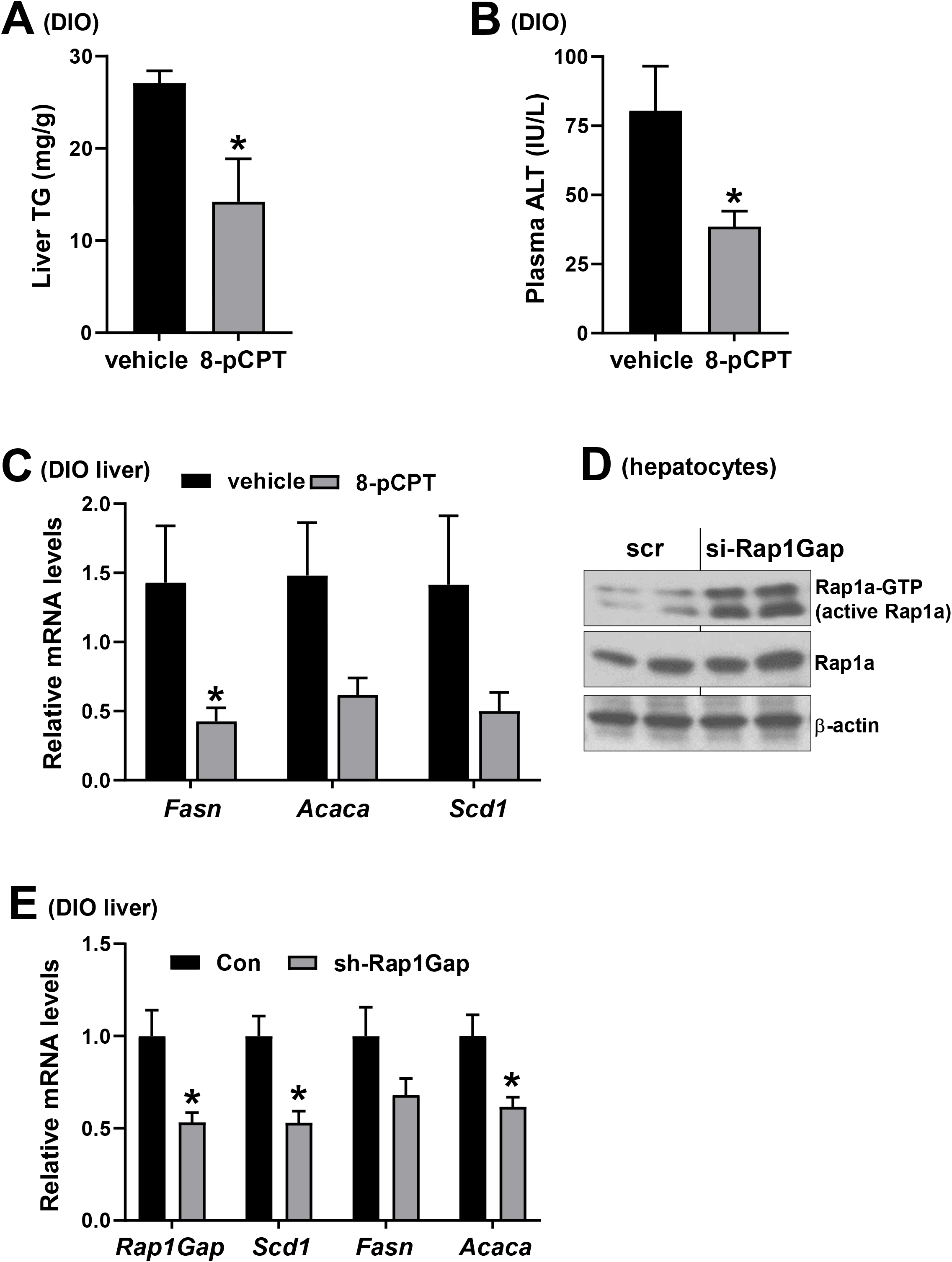
Rap1a activation via 8-pCPT or sh-Rap1Gap treatments lowers steatosis in DIO mice. (**A-C**) 20-week-old male DIO mice were treated with 1.5 mg/kg/day with 8-pCPT for 2 weeks. Hepatic triglyceride (TG) content (A), plasma ALT (B), and liver *Fasn, Acaca*, and *Scd1* mRNA levels (C) were assayed (n= 4 mice/ group, *p < 0.05). (**D**) GTP-Rap1a (active Rap1a) levels from cells treated with scr vs si-Rap1Gap. (**E**) 20-week-old male DIO mice were treated with an AAV-shRNA against Rap1Gap or control AAV and hepatic expression of Rap1Gap and lipogenic genes were analyzed (n= 10 mice/group; *p < 0.05).

Rap1Gap functions as a negative endogenous regulator of Rap1a by facilitating the hydrolysis of its GTP to GDP.^19^ We then investigated whether silencing Rap1Gap (si-Rap1Gap) can activate Rap1a and lower obesity-associated fatty liver formation. We first showed that si-Rap1Gap treatment increased Rap1a activity in primary hepatocytes (**Figure 2D**). Consistent with an increase in Rap1a activity, we found that the mice had lower expression of hepatic lipogenic gene expression (**Figure 2E**) when liver Rap1Gap was acutely silenced using AAV-H1-sh-Rap1Gap, which specifically deletes genes in hepatocytes.^20,21^ Altogether, these results suggest the therapeutic potential of Rap1a activators in protection against liver TG accumulation.

### Hepatic Rap1a regulates mature Srebp1 and mTORC1

As the genes involved in lipogenesis but not lipid oxidation were altered upon Rap1a activation, we next determined the unprocessed (precursor) and cleaved (mature) protein levels of master lipogenic transcription factor, Srebp1. We found that L-Rap1 KO DIO mice had increased mature, nuclear form of Srebp1 (**Figure 3A**). However, we observed that the unprocessed (precursor) form of Srebp1 was unchanged (**Figure 3A**). We obtained similar results in isolated hepatocytes treated with scr (control) vs siRNA against Rap1a (si-Rap1a) (**Figure 3B**). Conversely, TBG-CA-Rap1a treated DIO mice had lower mature form of Srebp1 without a change in the precursor form (**Figure 3C**). These data indicate that activation of Rap1a reduces obesity-associated hepatic liver TG accumulation, lipogenic gene expression, and cleavage of Srebp1, whereas Rap1a inhibition has the opposite effects.

**Figure 3.**
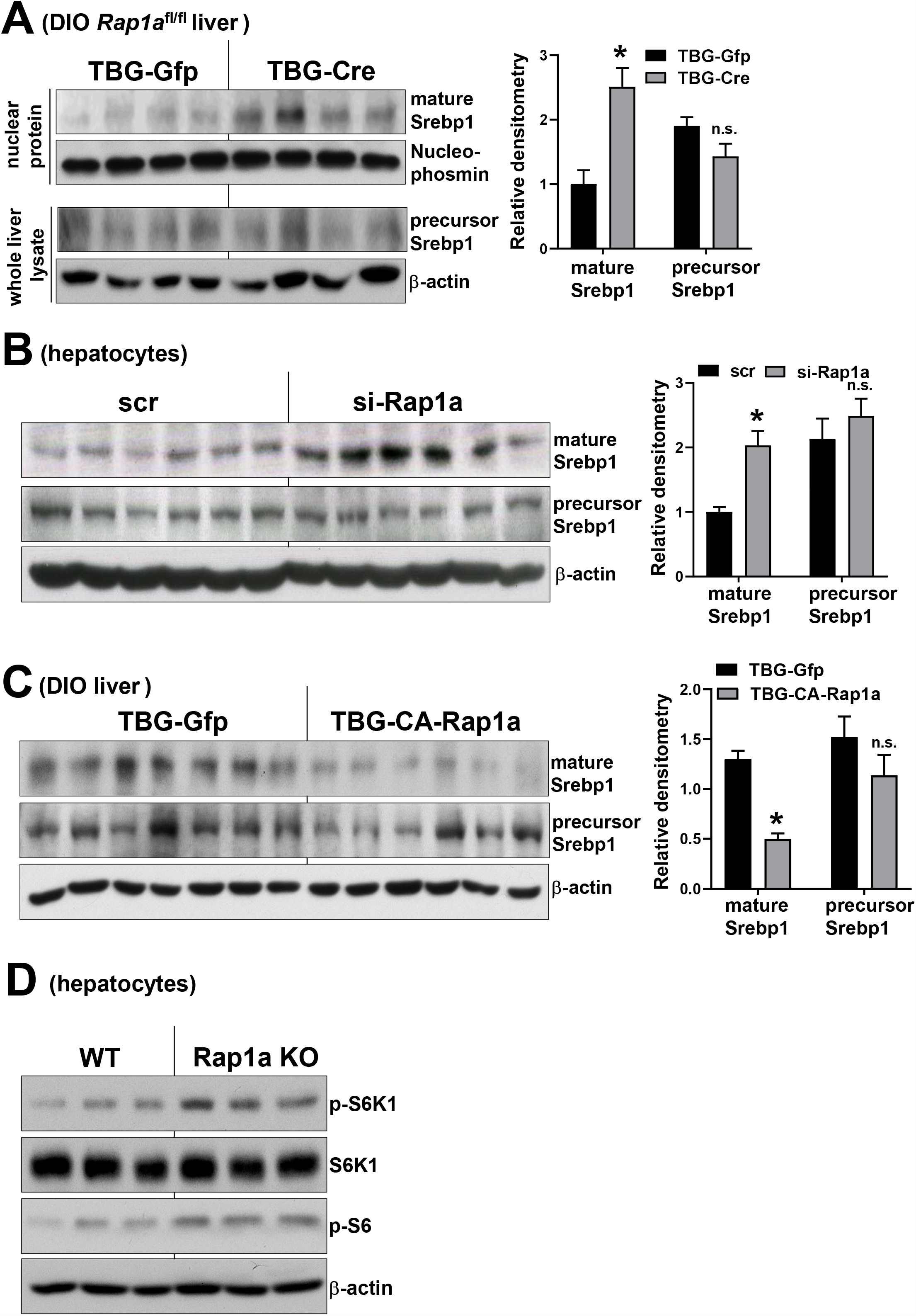
Hepatic Rap1a regulates mature Srebp1 and mTORC1. (**A-C**) Hepatic mature Srebp1 or precursor Srebp1 protein levels were assayed in the livers of *Rap1a*^fl/fl^ DIO mice treated with TBG-Cre or TBG-Gfp (control) (A), in DIO mice treated with TBG-Gfp or TBG-CA-Rap1a (B), or in scrambled (scr) or si-Rap1a treated hepatocytes (C). Densitometric quantification of the immunoblot data is shown in the graph. (**D**) p-S6K1 and p-S6 levels were analyzed in WT or Rap1a KO primary hepatocytes.

Mice lacking hepatic insulin receptor or both isoforms of Akt show defects in Srebp1 expression after feeding.^22,23^ This insulin induced Srebp1 upregulation is suggested to be mediated via mTORC1.^24,25^ Although the role of mTORC1 in the regulation of NAFLD and NASH is complex,^26-28^ its ability to regulate Srebp1 cleavage has been demonstrated in multiple mouse models.^29-31^ As inhibition of Rap1a suppresses insulin signaling and increases HGP,^12^ it might be predicted that Rap1a inhibition decreases mTORC1 activity. Surprisingly, we observed that mTORC1 is hyperactivated upon Rap1a deficiency as shown by increased phospho-S6K1 and phospho-S6 levels in isolated Rap1a KO hepatocytes as compared to controls (**Figure 3D**). This result is consistent with a recent study showing that Rap1a deficient cells have constitutive mTORC1 activity irrespective of insulin signaling.^32^

### Rap1a activity is decreased in NASH liver and restoring it lowers plasma ALT and liver fibrogenic gene expression

Steatosis is an important contributor to NASH pathogenesis and 20% of those individuals with NAFLD progress to NASH^9^, and thus, we next asked whether a causal relationship exists between hepatic Rap1a and NASH. We obtained human liver specimens of obese individuals with normal or NASH histology and found that Rap1a activity was significantly lower in liver extracts from subjects with NASH vs control as shown by lower GTP-bound (active) Rap1a levels (**Figure 4A**). In preparation for causation studies, we next explored Rap1a activity in mice fed with a NASH provoking high-fructose, <u>p</u>almitate, and <u>c</u>holesterol-rich diet (FPC). Mice fed with the FPC diet develop liver steatosis, inflammation, and fibrosis by 16 weeks of feeding, and FPC-fed mice livers show human-like NASH pathologic and molecular features.^21,33,34^ Similar to human data, we found lower hepatic Rap1a activity in 16 week-FPC diet fed mice as compared to chow (**Figure 4B**). To test the functional importance of Rap1a in NASH, we placed WT mice on FPC diet for 16 weeks and injected them with AAV8-TBG-CA-Rap1a or AAV8-TBG-Gfp at the same time as the diet initiation. This treatment, which increased hepatic Rap1a activity without altering body weight or food intake (not shown), significantly lowered plasma ALT (**Figure 4C**) and liver fibrogenic gene expression (**Figure 4D**).

**Figure 4.**
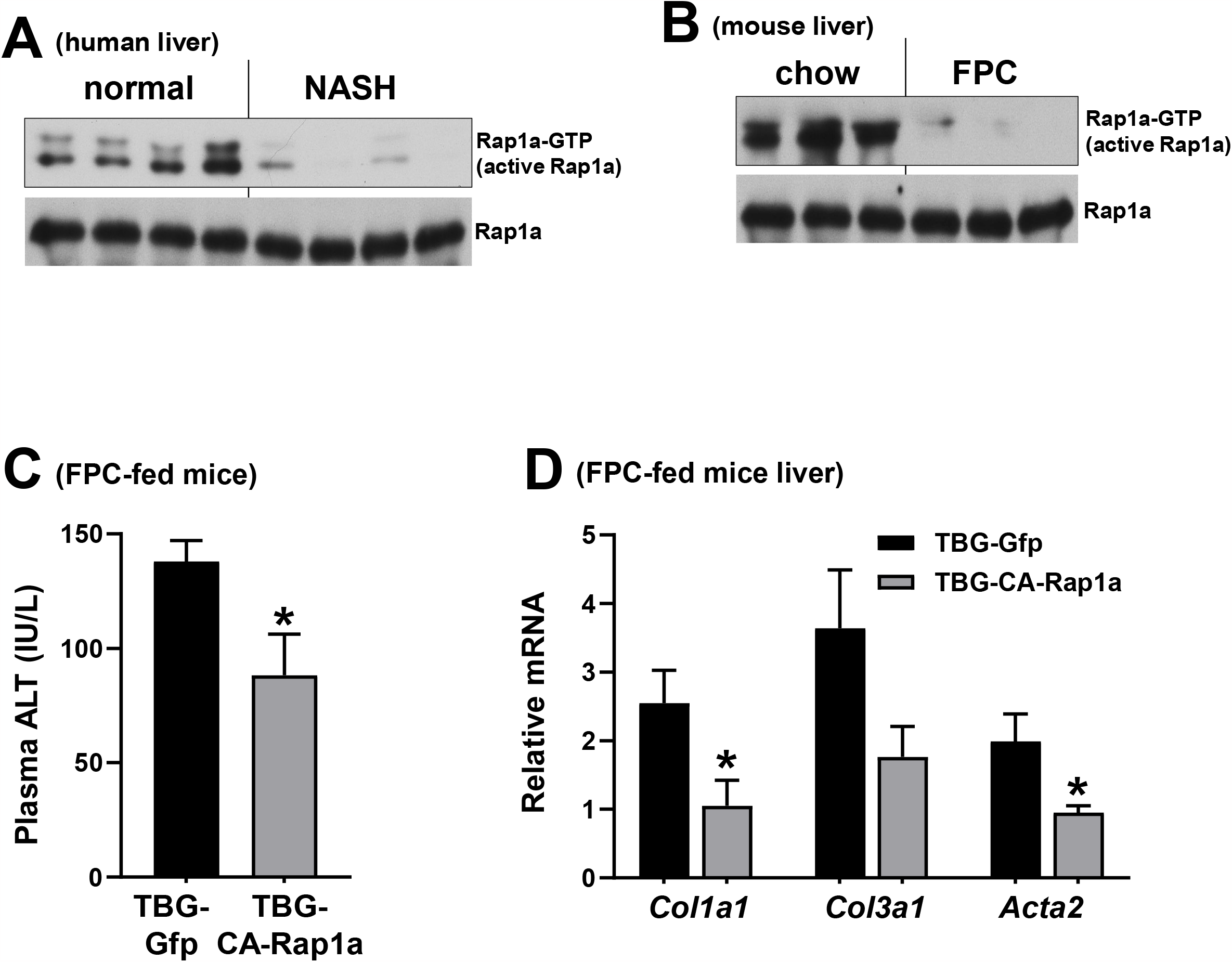
Rap1a activity is decreased in NASH liver and restoring it lowers plasma ALT and liver fibrogenic gene expression. (**A-B**) Rap1a-GTP (active Rap1a) levels were assayed in normal and NASH human liver (A) and chow vs FPC diet fed mice liver (B), and chow vs high fat-CDAA diet fed mice liver (HFD-CDAA) (C). (**C-D**) Plasma ALT (C) and hepatic fibrogenic gene mRNAs (D) were assayed in FPC-diet fed WT mice treated with AAV8-TBG-CA-Rap1a or AAV8-TBG-Gfp (control) (n= 4-5 mice/ group, *p < 0.05).

## Discussion

We previously reported that activation of hepatic Rap1a suppresses gluconeogenic gene expression and improves glucose intolerance via Akt mediated FoxO1 inhibition, suggesting that Rap1a activation enhances insulin signaling.^12^ Our results here demonstrate that, despite enhanced insulin action, Rap1a activation also protects against hepatosteatosis. The mechanism involves suppression of Srebp1 cleavage through mTORC1 inhibition, suggesting that the effects of Rap1a on insulin action can be uncoupled from the mechanisms regulating fatty acid metabolism. In the future, it would be interesting to investigate the molecular mechanisms by which Rap1a promotes insulin response but inhibits mTORC1. In this context, recent work has shown that Rap1a depletion results in hyperactivation of mTORC1 activity via expanding the lysosome population, which markedly increases the association between mTORC1 and its lysosome-borne activators.^32^

Statins inhibit Rap1a, which contributes to their hyperglycemic effect.^12^ As Rap1a inhibition results in increased liver TG accumulation, statins may be predicted to increase hepatic lipid content. Although mild transaminase elevation has been reported in patients with statin use,^35^ clinical trials assessing the effect of statins on NAFLD demonstrated mixed results for markers of inflammation and liver damage, and consistent histological data are still pending.^36^ It is important to note that in addition to inhibiting geranylgeranyl isoprenoids and Rap1a, statins have many other effects and could have important functions in other cell types that may have protective roles in the pathogenesis of NAFLD.^37^ Nonetheless, future work is needed to thoroughly investigate the effects of statins on NAFLD/NASH pathogenesis.

The results here and our previous work thus far suggested that activation of Rap1a in the liver is beneficial in obesity-associated cardiometabolic diseases as it i) lowers proatherogenic proprotein convertase subtilisin/kexin type 9 (PCSK9) and low-density lipoprotein cholesterol; ii) improves glucose intolerance; and iii) protects against NAFLD.^12,18^ Thus, therapeutically activating Rap1a could confer dual benefit in cardiometabolic disease, i.e., by both improving T2D/fatty liver formation and, through lowering plasma PCSK9 and LDL-C, protect against atherosclerosis. Moreover, as inhibiting Rap1Gap activates Rap1a and lowers HGP and hepatic lipogenic gene expression, silencing Rap1Gap could be another therapeutic approach to activate Rap1a and provide metabolic benefit. It is important to note that liver targeting N-acetylgalactosamine (GalNAc)-siRNA conjugates and similar hepatocyte-targeted siRNA approaches are FDA-approved for human use.^38-40^

## Materials and Methods

### Mouse Experiments

WT (stock # 000664), *db/db* and *db/+* (stock # 000697) and diet-induced obese (DIO, stock # 380050) mice were from Jackson Labs. DIO mice were fed a high-fat diet with 60% kcal from fat (Research Diets). NASH provoking high-fructose, <u>p</u>almitate, and <u>c</u>holesterol-rich diet (FPC) diet was from Envigo. *Rap1a*^fl/fl^ mice were generated by crossbreeding Rap1ab double floxed mice (Jackson Labs, stock number: 021066) with WT C57BL/6J mice and confirmed by genotyping. To avoid the effects of gender-related differences in metabolism, male mice were used for most of the experiments. All mice were maintained on a 12 h-light-dark cycle. Recombinant adeno-associated viruses (AAVs, 1-2 X 10^9^ genome copies per mouse) were delivered to mice by tail vein injections, and experiments were commenced after 3-28 days. 20-week-old male DIO mice were treated intraperitoneally with 8-pCPT 1.5 mg/kg/day for 2 weeks. For all experiments, male mice of the same age and similar weight were randomly assigned to experimental and control groups. Animal studies were conducted in accordance with the Columbia University Irving Medical Center Animal Care and Use Committee.

### Human liver samples

Human liver samples were obtained from the NIH-supported Liver Tissue Cell Distribution System at the University of Minnesota. The samples were collected postmortem on the date of liver transplantation and preserved as frozen samples. The Institutional Review Board at the Columbia University Medical Center approved the research protocol.

### Reagents and Antibodies

Anti-β-actin, anti-p-S6K1 and anti-p-S6 antibodies were from Cell Signaling Technology. Anti-Rap1a antibody (cat # AF3767) was from R&D Systems. AAV8-TBG-Cre and AAV8-TBG-Gfp were obtained from Gene Therapy Resource Program (GTRP) of NHLBI or purchased from Addgene (cat # 105535-AAV8 and # 107787-AAV8). AAV8-TBG-CA-Rap1a was from Penn Vector Core. Specific Epac activator, 8-pCPT, was from Cayman Chemicals (cat # 17143). AAV8-shRNA targeting murine *Rap1Gap* was made by annealing complementary oligonucleotides, which were then ligated into the self-complementary AAV8-RSV-GFP-H1 vector as described previously.^41^ The resultant construct were amplified by Salk Institute Gene Transfer, Targeting, and Therapeutics Core.

### Liver Triglyceride Measurement

Liver triglyceride measurement was done as described previously.^42^ Briefly, the lipids were extracted using a modification of the Bligh-Dyer method (Bligh and Dyer, 1959). Livers were homogenized in chloroform: MeOH: H2O (1:2:0.8) at room temperature and then centrifuged. Equal volumes of chloroform and water were added to the supernatant fraction, which was then vortexed and centrifuged. The chloroform layer was collected and dried under nitrogen. The dried lipids were then resuspended in 90% isopropanol: 10% Triton-X and then assayed for triglyceride using a kit from Wako and cholesterol using a kit from Life Technologies.

### Hepatocyte experiments

Primary mouse hepatocytes were isolated from 12-20-week-old male or female WT mice as described previously.^43^ Transfections with scrambled or si-Rap1a, constructs were carried out using Lipofectamine RNAiMAX reagent according to manufacturer’s instructions.

### Immunoblotting

Liver protein was extracted using RIPA buffer (Thermo Scientific), and the protein concentration was measured by DC protein assay kit (Bio-Rad). Whole cell lysates were harvested in 2X Laemmli buffer. Cell extracts were electrophoresed on SDS-polyacrylamide gels and transferred to 0.2 μm or 0.45 μm PVDF membranes. Blots were blocked in Tris-buffered saline with 0.1% Tween-20 containing 5% BSA at room temperature for one hour. Membranes were then incubated overnight at 4°C with primary antibodies. The protein bands were detected with horseradish peroxidase-conjugated secondary antibodies (Jackson ImmunoResearch) and Supersignal West Pico enhanced chemiluminescent solution (Thermo).

### Rap1a Activity Assay

Rap1a activity was assayed using Active Rap1 Detection Kit (Cell Signaling Technology, cat # 8818). Liver tissue sample were lysed with lysis/binding/washing buffer containing 2 mM PMSF, 5μg/ml leupeptin, 10 nM okadaic acid and 5μg/ml aprotinin. GST-RalGDS-RBD fusion protein was used to bind the activated form of GTP-bound Rap1a, which was then immunoprecipitated with glutathione resin. Rap1a activation levels were determined by western blot using a Rap1a primary antibody.

### Quantitative PCR

RNA was isolated from hepatocytes or liver tissue using TRIzol (Invitrogen) and cDNA was synthesized from 1 ug of RNA using a cDNA synthesis kit (Invitrogen). Real-time PCR was performed using a 7500 Real-Time PCR system and SYBR Green reagents (Applied Biosystems).

### Statistical analysis

All results are presented as mean ± SEM. *P* values were calculated using the Student’s *t*-test for normally distributed data and the Mann-Whitney rank sum test for non-normally distributed data. For experiments with more than two groups, *P* values were calculated using one-way ANOVA for normally distributed data and the Kruskal-Wallis by ranks test for non-normally distributed data.

